# Smith-Lemli-Opitz syndrome carrier frequency and estimates of *in utero* mortality rates

**DOI:** 10.1101/085084

**Authors:** Gabriel A. Lazarin, Imran S. Haque, Eric A. Evans, James D. Goldberg

## Abstract

Objective: To tabulate individual allele frequencies and total carrier frequency for Smith-Lemli-Opitz syndrome (SLOS) and compare expected versus observed birth incidences.

Design: Retrospective analysis of patients from the general population who have undergone carrier screening for SLOS.

Setting: Individuals were offered and elected carrier screening in their respective physician’s offices.

Patients: 262,399 individuals with no indication of family or personal history of Smith-Lemli-Opitz syndrome, primarily US-based, screened for Smith-Lemli-Opitz syndrome mutations as part of an expanded carrier screening panel.

Intervention(s): Data on mutations in the *DHCR7* gene causing SLOS were analyzed to estimate carrier frequencies in multiple ethnic groups. SLOS birth incidences obtained from existing literature are then compared to our data to estimate the effect of SLOS on fetal survival.

Main Outcome Measure(s): Individual and cumulative allele frequencies stratified by self-reported patient ethnicity.

Results: SLOS carrier frequency is highest in individuals of Ashkenazi Jewish ancestry (1 in 43) and Northern Europeans (1 in 54). Comparing predicted birth incidence to that observed in published literature suggests that approximately 42% to 88% of affected conceptuses experience fetal demise.

Conclusion: SLOS is relatively frequent in certain populations and, due to its impact on fetal survival, merits preconception screening.

## BACKGROUND

Smith-Lemli-Opitz syndrome (SLOS, OMIM #270400) is an autosomal recessive disease caused by mutations in the *DHCR7* gene. Individuals with the disease exhibit a wide and variable spectrum of phenotypic abnormalities, including multiple congenital malformations, facial abnormalities, metabolic errors, and intellectual disability. Fetal demise may be a relatively common outcome, occurring in up to 80% of affected fetuses (1). Table 1 lists all characteristics that may occur. Variable, and sometimes subtle, presentation can lead to missed or delayed diagnoses (2). Prenatally, non-specific ultrasound findings may be present, such as cardiac defects or cleft lip/palate. Prenatal biochemical screening approaches (4) are also available.

Disease frequency estimates have varied due to methods of ascertainment, alleles assessed, and populations studied. In general, existing data suggest a carrier frequency of approximately 1% for common alleles in Caucasians (5, 6, 7, 16), with at least one source extrapolating the total carrier frequency to 3% (8). The most common allele in North American populations is the null mutation, c.964-1G>C, while other alleles, c.452G>A and c.278C>T, may be more frequent in Central European and Mediterranean ancestry populations, respectively (9).

SLOS disease incidence has been studied, primarily in Europe and Canada. Diagnoses have been confirmed by molecular and biochemical methods. Most figures range from 1/60,000 (10, 11) to 1/20,000 (12, 13, 14). A large study of SLOS risk assessed in over a million pregnancies in the US found a mid-trimester SLOS prevalence of 1/101,000 Caucasians, much lower than other estimates (4). Elevated risk was initially identified by mid-trimester serum analysis. However, since SLOS diagnostic testing was not performed in a number of screen-positive pregnancies (in particular those with fetal demise), this data underestimates the incidence at conception when SLOS results in first trimester or embryonic lethality. The authors do not comment on possible reasons for the discrepancy between their findings and those of other population studies.

Data regarding other ethnic populations are limited, but where available, suggest that SLOS is uncommon or rare in non-Caucasians, particularly among individuals of African or East Asian ancestry (6, 7, 15, 16).

This study utilizes a large database of individuals tested for SLOS to report observed carrier frequencies and estimate the expected birth incidence resulting from those frequencies. 262,399 individuals with no reported indication of personal or family history of SLOS or infertility were screened for SLOS mutations as part of an expanded carrier screening panel, including samples of more than 10,000 for most major US ethnic groups. Because this population is large and screened without apparent indication or dependency on clinical symptoms, highly accurate allele frequency estimates are possible.

## METHODS

This is a retrospective analysis of results from individuals electing expanded carrier screening that included Smith-Lemli-Opitz syndrome between January 2012 and December 2015.

The analyses for this study were performed in a CLIA and CAP-certified laboratory using two methods (Family Prep Screen 1.0 and 2.0, Counsyl, South San Francisco, CA). Most (n=210,857) were screened via targeted genotyping (Family Prep Screen 1.0) for 13 *DHCR7* mutations using TaqMan fluorescent probes on the Fluidigm 96.96 platform. These mutations were included in the original study referenced in the Introduction. The original panel in addition had four more mutations that were since removed due to low prevalence.

Another 51,542 were screened via a next-generation sequencing test (Family Prep Screen 2.0) using custom hybrid capture followed by sequencing on the Illumina HiSeq 2500 to test for variants in *DHCR7* exons 3-9. This methodology encompasses the 13 mutations identified by genotyping, the additional four included in the original study, and other mutations previously known or undescribed. Large deletions and insertions, which may account for 4-5% of causative alleles (9, 17), would typically not be identified from this methodology. Identified variants were classified for pathogenicity based on the American College of Medical Genetics and Genomics' recommendations for interpretation and reporting using the approach described by Karimi, *et al* (18). Patients were informed when a known, likely or predicted deleterious variant was identified. The combination of test methodology, variant classification and variant reporting will be referred heretofore as NGS. Variants of uncertain significance and known, likely or predicted benign variants were not routinely reported to the physician or patients, as per our laboratory’s routine carrier screening protocol.

### Study Population

This population totals 262,399 individuals that elected expanded carrier screening that included SLOS between January 2012 and December 2015. Carrier status for up to 109 genes in addition to *DHCR7* could be assessed simultaneously. The laboratory’s total tested population within this time range is greater than 262,399, but individuals were excluded from this analysis when any of the following occurred: an indication other than “no family history (routine carrier screening)” was selected, SLOS was not included in a customized disease panel ordered by the physician, or the patient requested exclusion of his/her results for research purposes.

The ordering physician or the patient directly reported ethnicity. Unknown ethnicity could be selected. These unknown individuals and ones for which no response was selected are reported together.

All tests were ordered by a physician or other health care provider. Most were obstetricians, maternal fetal medicine specialists, reproductive endocrinologists, geneticists and genetic counselors. Institutional review board exemption is applicable due to de-identification of the data presented (45 CFR part 46.101(b)(4)). Follow-up genetic counseling was made available at no cost to all individuals tested. Testing was performed as fee-for-service, typically paid for by a third-party and/or the patient.

## RESULTS

Data for ethnicities where *n* > 9,000 and carrier frequency exceeds 0.5% are detailed in Table 2. The supplementary section includes the remaining populations.

### Patient demographics

Of 210,857 that had the genotyping assay, Mixed / Other Caucasians represented the largest reported ethnic group (25.14%) followed by Northern Europeans (23.40%). Finnish represented the smallest ethnic group (0.07%) and Native Americans were the smallest of the major US ethnic groups (0.18%). Nearly 14% of the tested population had unknown or unreported ethnicity.

### Targeted mutation data

Of 10 ethnic groups with *n* > 3000, the highest carrier frequency was found among Ashkenazi Jewish (2.35% or 1/42) and the lowest among South Asians (0.07% or 1/1477). In general, the frequency was low among Asian populations. On the other hand, all populations of European origin showed carrier frequencies exceeding 1%.

Of the 13 targeted mutations assayed, all were detected six times at minimum and 11 of the mutations were detected at least 10 times. Nonetheless, two were predominantly frequent. The null c.964-1G>C mutation was most frequent, accounting for 75.0% of carriers identified. It was the most frequent, or tied for most frequent, mutation identified in non-Asian ethnic groups. But, these latter populations had few carriers identified. Where c.964-1G>C was the most frequent mutation, we observed varying carrier frequency, ranging from 2.14% in Ashkenazi Jewish to 0.10% in Middle Easterners

The second most frequent allele was c.452G>A, accounting for 16.5% of all carriers’ mutations. It was most common in the Cajun/French-Canadian population, with a carrier frequency of 0.52%.

### Next-generation sequencing data

Included in the targeted mutation dataset above, 51,542 individuals underwent comprehensive mutation analysis through NGS. The same eligibility criteria apply to these data as described in the Methods.

The patient demographic pattern approximates that of the larger genotyped population. Mixed / Other Caucasians (25.4%) and Northern Europeans (17.7%) were the largest populations. Greater than 800 individuals were tested in 10 ethnic groups, ranging from 834 (Southeast Asian) to 13,073 (Mixed / Other Caucasian).

As expected, in most ethnic groups, the carrier frequency by comprehensive analysis was higher compared with that by targeted analysis. The relative increase varied. A greater increase was observed among non-Caucasian groups, which also had the lowest initial frequency. This is logical; the targeted panel was based off studies primarily conducted in European populations and even the most common alleles were infrequent among non-European groups. Therefore discovery of additional infrequent alleles would have greater impact on overall carrier tabulations.

Finally, in order to elucidate the benefit conferred by the NGS approach, the percentage of carriers identified by NGS and *not* identified by targeted analysis were calculated. This ranged from 0% (four ethnic groups) to 80% (East Asians), and overall the targeted approach detected 92.4% of all of the mutations detected in this predominantly European population (59% of individuals). Table 3 details, among only the population tested by NGS, the numbers of mutations that were included on the 13 mutation panel or the NGS panel.

**Table 3.** Comparison of NGS and TG methodologies for Smith-Lemli-Opitz syndrome carrier detection in selected populations (n>800).

| **Ethnicity** | **Tested by**<br>**NGS, *n*** | **All carriers detected by**<br>**NGS, *n*** | **Carriers detectable**<br>**by TG panel, *n*** | **Carriers missed by TG panel** |
| --- | --- | --- | --- | --- |
| **African** | 3,284 | 14 | 14 | 0% |
| **Ashkenazi Jewish** | 4,695 | 103 | 103 | 0% |
| **Mixed/Other Caucasian** | 13,073 | 258 | 240 | 7% |
| **East Asian** | 3,102 | 5 | 1 | 80% |
| **Hispanic** | 3,377 | 29 | 27 | 7% |
| **Middle Eastern** | 861 | 4 | 2 | 50% |
| **Northern European** | 9,109 | 178 | 165 | 7% |
| **South Asian** | 1,872 | 3 | 1 | 67% |
| **Southeast Asian** | 834 | 2 | 1 | 50% |
| **Southern European** | 1,512 | 31 | 28 | 10% |
| **Unknown** | 9,518 | 128 | 115 | 10% |

In total, the NGS approach identified 58 occurrences of 30 unique non-TG mutations. Three mutations were identified in more than three individuals; c.1337G>A was identified nine times in five patient populations.

One potentially “affected” individual was identified in the NGS dataset: a person that was compound heterozygous for two *DHCR7* mutations: c.111G>A and c.429T>G. The individual underwent genetic counseling and no related symptoms were apparently reported. Due to other present clinical circumstances, further investigation was not initiated at that time. Possible explanations include: unreported or unknown clinical symptoms or diagnosis, *cis* configuration of alleles, genetic “diagnosis” with other modifying/alleviating factor, or laboratory error.

### Impact on Fetal Survival Rates

Disease incidence estimates range from 1/101,000 to 1/20,000, per above. The largest non-mixed population, Northern Europeans (n=58,439), were commonly studied in those literature sources as well. SLOS birth incidence based on Hardy-Weinberg principles is predicted to be 1/11,664 based on the following calculation:

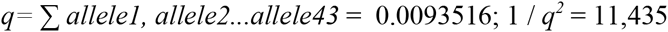

Using the lowest and highest birth incidence findings suggests an *in utero* demise rate of 88% to 42%.

## DISCUSSION

Accurate carrier frequencies for Smith-Lemli-Opitz syndrome are reported here, based on screening of a large general population cohort. Frequencies are approximately 2% in Caucasians and Ashkenazi Jewish and exceed 0.5% in Hispanics and African Americans. These are meaningful since current carrier screening guidelines included diseases of similar frequency (19). Comparisons of the disease’s predicted birth incidence from the data presented here and observed birth incidences from the literature suggest a significant proportion of affected conceptuses do not survive.

The overall carrier frequency for this population is 1.4%, though this again has limited application to an individual clinical setting, given substantial ethnic variability. SLOS carriers are most frequent among individuals of European ancestry, in particular Northern European and Ashkenazi Jewish. While previous disease incidence estimates have ranged from 1/20,000 to 1/101,000, these data predict an incidence at the higher end of that spectrum - at conception, 1/11,664 in Northern Europeans and 1/7,396 in Ashkenazi Jewish. Combining all Caucasian populations yields a carrier frequency of 1.7%, and a predicted disease incidence at conception of 1/13,924.

In Hispanics and African Americans, carrier frequencies are 1/167 and 1/183, respectively. In these populations, predicted disease incidences are approximately 1/111,556 to 1 in 133,956. Carrier status for SLOS is very rare among all Asian populations we studied.

Differences between birth observation rates and these predictions may be due to the significant *in utero* mortality rate. Fetal demise has previously been suggested to occur in up to 80% of fetuses affected with SLOS (1). Hydrops has been described in several cases of fetuses later diagnosed with SLOS, though it is also clear that this is not an inevitable outcome. It is noteworthy that a study in the Icelandic population predicted finding 19.1 individuals homozygous for c.964-1G>C in a population of 104,220 but actually found none, further suggesting early lethality of this genotype (20). Craig, *et al.*, reported a large study of over a million pregnancies biochemically screened for SLOS (4). They estimated a mid-trimester prevalence of 1/101,000 Caucasians. Two considerations in evaluating this difference are that 30% of SLOS screen-positive fetuses were excluded from analysis due to fetal demise and that the biochemical screening performed in the second trimester does not detect conditions with first trimester lethality. Continued research may provide explanation, but the data here, in combination with those of Craig, *et al*, suggest that first or second trimester demise are the most likely outcome of SLOS-affected fetuses. That likelihood depends on the true live birth incidence, but based on most estimates the prenatal mortality rate is 42-88%.

In the only other related study located that utilized NGS methodology, Cross, *et al*, examined the frequency of *DHCR7* pathogenic variants in the 1000 Genomes population (21). In that, they found a 1.01% carrier frequency and predicted a disease incidence of 1/39,215 conceptions. However, they pool a number of non-Northern European populations (Colombian, Iberian, Puerto Rican, Toscani) into their Northern European pool. The data here indicate that this pooling undercounts the actual frequency, since Hispanics and Southern Europeans have lower carrier frequencies. Restricting analysis to Northern European populations (British, Utah, Finnish) shows 6 of 290 (2.01%) individuals to be carriers for the c.964-1G>C variant alone.

This study’s foremost limitation is that ethnicity reporting is based on the patient or clinic’s report and may therefore be erroneously classified. In addition, the laboratory restricts selection to a single ethnic group - an unknown number of individuals have multiple ancestral backgrounds and these are not accounted for. Ascertainment is also incomplete, since an individual had to elect carrier screening to be included in the dataset. Bias in minimized by limiting the dataset to individuals that reported no predisposition toward positive SLOS carrier status, but this does not account for how the data may differ from the untested cohort. Lastly, neither test methodology routinely detected large copy number variants. A similar large-scale study inclusive of these variant types would help further define the full mutation spectrum.

These data present Smith-Lemli-Opitz syndrome carrier frequencies obtained from large-scale routine carrier screening and suggest a substantial *in utero* mortality rate. These are the largest sample sizes reported to date of every major US-based population. Given the relatively high carrier frequency in a subset of populations and the impact on fetal survival, preconception carrier screening is suggested.

## Tables

- Table 1: <u>List of SLO characteristics</u>
- Table 2: Carrier<u>frequencies for selected populations</u>
- Table 3: <u>TG vs NGS comparison</u>
- Supplementary table: Carrier<u>frequencies for additional populations</u>

## Smith-Lemli-Opitz syndrome phenotypic features

### General

*in utero* demise
prenatal and postnatal growth retardation
feeding problems (poor suck, irritability, failure to thrive)
behavioral problems (sensory hyperreactivity, irritability, sleep cycle disturbance, self-injurious behavior, autism spectrum behaviors, temperament dysregulation, social and communication defects, depression)

### Central Nervous System / Neurological

intellectual disability, mild to severe
hypotonia
seizures
abnormalities of myelination
ventricular dilatation
malformations of the corpus callosum and/or cerebellum
Dandy-Walker malformation or variants
holoprosencephaly

### Facial

characteristic facies (narrow forehead, epicanthal folds, ptosis, short nose with anteverted nares, short mandible with preservation of jaw width, capillary hemangioma over the nasal root, low set posteriorly rotated ears)
cleft lip/palate
abnormal gingivae

### Cardiac / Pulmonary

functional defects
single chamber or vessel defects
complex cardiac malformations
abnormal pulmonary lobation
pulmonary hypoplasia

### Genital

hypospadias
ambiguous genitalia (failure of masculinization of male genitalia)

### Skeletal

postaxial polydactyly
2-3 toe syndactyly
microcephaly

### Abdominal

gastrointestinal problems (dysmotility, hypomotility, gastrointestinal reflux, constipation, formula intolerance)
liver disease
renal hypoplasia or agenesis
renal cortical cysts
hydronephrosis
structural anomalies of the collecting system
splenomegaly

### Ocular

photosensitivity
congenital cataracts,
strabismus
optic atrophy
optic nerve hypoplasia

### Endocrine

electrolyte abnormalities
hypoglycemia
hypertension
adrenal insufficiency
low testosterone in males

### Ear

otitis media
hearing loss

Based on Kelley (2000), GeneReviews.

|  |  |  |  |  |  |  |  |
| --- | --- | --- | --- | --- | --- | --- | --- |
| c.278C>T | T93M | 2 | 0 | 8 | 7 | 7 | 1 |
| c.452G>A | W151* | 5 | 37 | 229 | 11 | 178 | 40 |
| c.506C>T | S169L | 0 | 0 | 2 | 1 | 3 | 1 |
| c.724C>T | R242C | 2 | 0 | 28 | 0 | 27 | 1 |
| c.725G>A | R242H | 1 | 0 | 8 | 2 | 6 | 2 |
| c.906C>G | F302L | 0 | 0 | 0 | 3 | 0 | 0 |
| c.964-1G>C | IVS8-1G>C | 59 | 410 | 866 | 90 | 811 | 85 |
| c.976G>T | V326L | 0 | 3 | 11 | 0 | 15 | 0 |
| **Tested by NGS, with NGS-specific alleles below** |  | **3,284** | **4,695** | **13,073** | **3,377** | **9,109** | **1,512** |
| c.964-1G>T |  | 0 | 0 | 1 | 0 | 0 | 0 |
| c.1057delG | aka V353Wfs*60 | 0 | 0 | 0 | 0 | 1 | 0 |
| c.1139G>A | C380Y | 0 | 0 | 1 | 0 | 0 | 0 |
| c.1222T>C | Y408H | 0 | 0 | 1 | 0 | 0 | 0 |
| c.1295A>G | Y432C | 0 | 0 | 1 | 0 | 0 | 0 |
| c.1337G>A | R446Q | 0 | 0 | 3 | 1 | 2 | 1 |
| c.1389insT |  | 0 | 0 | 1 | 0 | 0 | 0 |
| c.1426T>C | aka<br>p.*476Qext*51 | 0 | 0 | 1 | 0 | 0 | 0 |
| c.1A>G | M1V | 0 | 0 | 0 | 0 | 1 | 1 |
| c.292C>T | Q98* | 0 | 0 | 1 | 0 | 0 | 0 |
| c.355delC | aka p.H119Ifs*8 | 0 | 0 | 0 | 0 | 1 | 0 |
| c.3G>A | M1I | 0 | 0 | 1 | 0 | 0 | 0 |
| c.413-2A>G |  | 0 | 0 | 0 | 0 | 1 | 0 |
| c.461C>G | T154R | 0 | 0 | 3 | 0 | 3 | 0 |
| c.461C>T | T154M | 0 | 0 | 1 | 0 | 0 | 0 |
| c.546G>A | W182* | 0 | 0 | 0 | 0 | 1 | 1 |
| c.651C>A | Y217* | 0 | 0 | 0 | 1 | 0 | 0 |
| c.952delT |  | 0 | 0 | 0 | 0 | 1 | 0 |
| c.963+1G>A |  | 0 | 0 | 1 | 0 | 0 | 0 |
| c.964-1G>T |  | 0 | 0 | 2 | 0 | 2 | 0 |
| **Cumulative Frequency (TG and NGS)** |  | **76 (0.55%)** | **452 (2.32%)** | **1,207 (1.83%)** | **121 (0.60%)** | **1093 (1.87%)** | **143 (1.51%)** |
|  |  | **1 in 183** | **1 in 43** | **1 in 55** | **1 in 167** | **1 in 54** | **1 in 66** |

